# Mitochondrial ATP synthase dimers spontaneously self-associate driven by a long-ranged membrane-induced force

**DOI:** 10.1101/272146

**Authors:** Claudio Anselmi, Karen M. Davies, José D. Faraldo-Gómez

## Abstract

ATP synthases populate the inner membranes of mitochondria, where they produce the majority of the ATP required by the cell. Cryo-electron tomograms of these membranes from yeast to vertebrates have consistently revealed a very precise organization of these enzymes. Rather than being scattered throughout the membrane, the ATP synthases form dimers, and these dimers are organized into rows that extend for hundreds of nanometers. These rows are only observed in the membrane invaginations known as cristae, specifically along their sharply curved edges. Although the presence of these macromolecular structures has been irrefutably linked to the proper development of cristae morphology, it has been unclear what drives the formation of the rows and why they are specifically localized in the cristae. We present the result of a quantitative molecular-simulation analysis that strongly suggests that the ATP synthase dimers organize into rows spontaneously, driven by a long-ranged attractive force that results from relief in the overall elastic strain of the membrane. This strain is caused by the V-like shape of the dimers, unique among membrane-protein complexes, which induces a strong deformation in the surrounding membrane. The process of row formation is therefore not a result of protein-protein interactions, or of a specific lipid composition of the membrane. We further hypothesize that once assembled, the ATP synthase dimer rows prime the inner mitochondrial membrane to develop folds and invaginations, by causing macroscopic membrane ridges that ultimately become the cristae edges. In this view, mitochondrial ATP synthases would contribute to the generation of a morphology that maximizes the surface area of the inner membrane, and thus ATP production. Finally, we outline the key experiments that would be required to verify or refute this hypothesis.

## Introduction

ATP synthases play a central role in cellular bioenergetics. Located in the plasma membrane of bacteria, archaea and cyanobacteria, or the innermost membranes of chloroplasts and mitochondria, these ubiquitous enzymes harness transmembrane electrochemical gradients of H^+^ or Na^+^, generated from respiration or light harvesting, to produce the majority of ATP used by the cell.

ATP synthases consist of four functionally distinct elements: a transmembrane domain, a catalytic domain, and a central and peripheral stalk. H^+^ or Na^+^ permeating through the transmembrane region, down their electrochemical gradients, cause the rotation of a sub-complex known as the c-ring. This rotation is transmitted to the catalytic region via the central stalk, which elicits a cycle of conformational changes that result in the synthesis and release of ATP. These changes in the internal structure of the catalytic domain are possible because the peripheral stalk holds the catalytic region stationary relative to the transmembrane region. Thus, the exergonic process of ion permeation is efficiently coupled to the endergonic process of ATP synthesis.

Although the architecture of the ATP synthase complex is largely conserved across species, the mitochondrial enzyme is distinct in that it exists as a dimer. This dimer is formed through accessory transmembrane subunits not present in prokaryotes or chloroplasts (Arnold et al., 1998; Lapaille et al., 2010). The dimers of mitochondrial ATP synthases are unique compared to other membrane-protein oligomers in that the two transmembrane domains are tilted relative to each other, approximately at a right angle (Allegretti et al., 2015; Hahn et al., 2016; Guo et al., 2017; Klusch et al., 2017). This arrangement causes the ATP synthase dimer to have a striking V-like shape, with the two catalytic domains projecting away from one another. In addition, these complexes are also remarkable in that they are not scattered randomly throughout the inner mitochondrial membrane; instead, they are organized into long rows that run along the sharply curved edges of the membrane invaginations called cristae (Strauss et al., 2008; Dudkina et al., 2010; Davies et al., 2011; Davies et al., 2012; Muhleip et al., 2017). These rows extend for hundreds of nanometers and include dozens of dimers, arranged side-by-side (**Fig. 1**).

**Figure 1.**
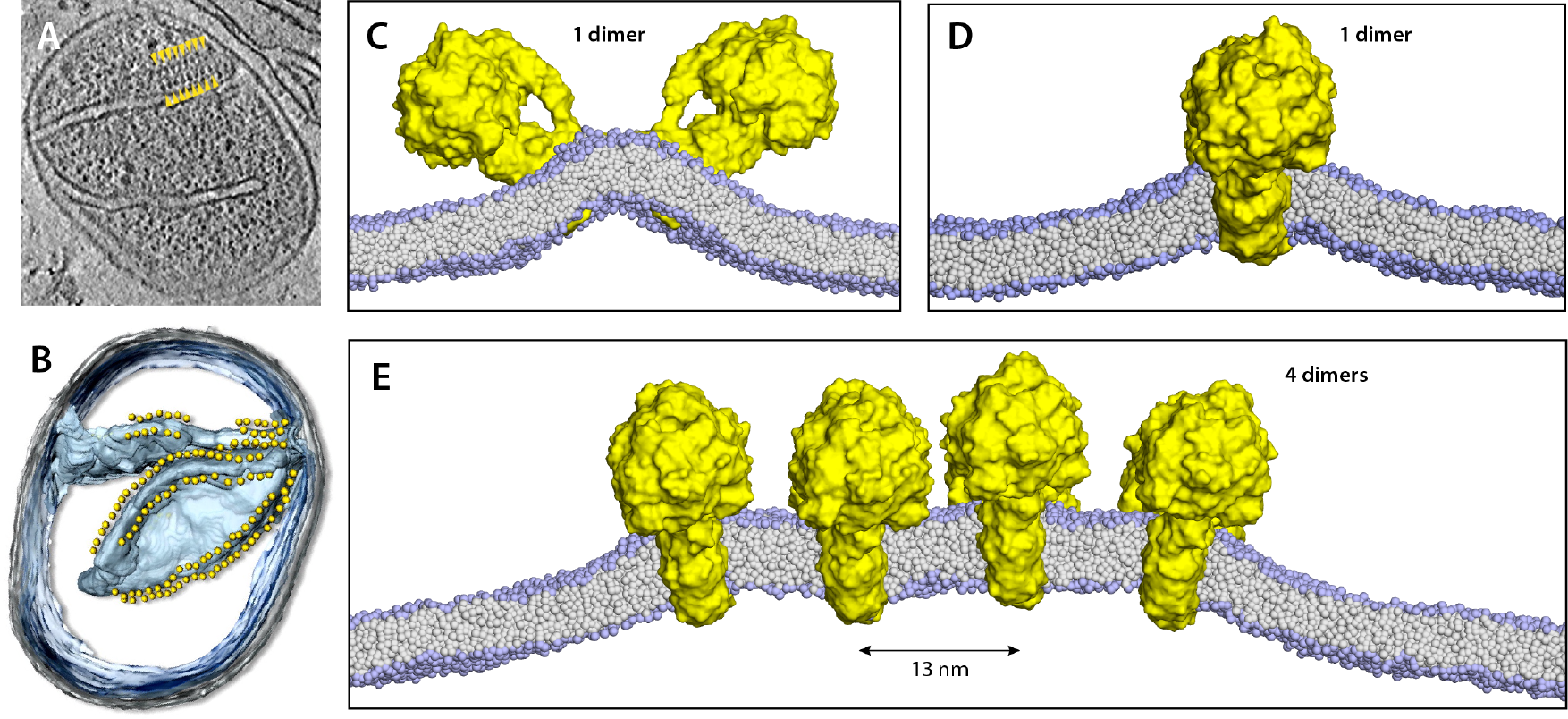
Organization of mitochondrial ATP synthase dimers and their membrane-bending properties. Panels (A-B) are adapted from (Davies et al., 2011); Panels (C-D) from (Davies et al., 2012). (**A**) Tomographic slice through an intact mitochondrion of *Podospora anserina*; yellow arrow heads mark the location of the ATP synthase dimers. (**B**) Three-dimensional reconstruction of the data in (A); yellow spheres indicate the catalytic domains in the ATP synthase dimers, against the cristae membrane (cyan). (**C**) Curvature perturbation caused by an isolated ATP synthase dimer along the direction parallel to the dimer long axis, from coarse-grained molecular dynamics simulations. (**D**) Same as C, along the direction perpendicular to the dimer. (**E**) Same as (D), for a row of four ATP synthase dimers, arranged side-by-side as observed in mitochondrial cristae. Note that the membrane is flat in between adjacent dimers along the direction of the row. In the perpendicular direction, the membrane profile is nearly identical to that in (C).

The factors that drive and stabilize these supramolecular structures have been unclear. Using molecular dynamics simulations, we have previously shown that an isolated ATP synthase dimer induces a pronounced, long-ranged deformation in the surrounding membrane, owing to its V-like shape (**Fig. 1**) (Davies et al., 2012). This curvature deformation occurs along both the direction parallel and perpendicular to the dimer interface. By contrast, when multiple dimers were placed side-by-side and kept a distance similar to that observed in mitochondria, we observed that the membrane flattens in between adjacent dimers, i.e. it restores its natural shape along the direction of the row. Based on a simple elastic model (Helfrich, 1973; Marsh, 2006), we *inferred* from these different curvatures that the membrane might drive the formation of the ATP synthase dimer rows, to reduce its overall deformation (Davies et al., 2012). However, direct evidence that this effective force in fact *emerges* from the interatomic interactions and dynamics of the molecular system was not obtained. Here, we present free-energy molecular simulations designed to explicitly probe the existence and magnitude of this hypothetical membrane-induced attractive force. We also discuss the implications of our findings and how to test them experimentally.

## Methods

### Molecular systems and simulation parameters

Two molecular systems were used in this study, both based on the MARTINI 2.1 force field (Marrink et al., 2007). The first system, or system A, consists of a bilayer of 5,832 1-palmitoyl-2-oleoyl-sn-glycero-3-phospho-choline (POPC) lipids, in a 100 mM NaCl solution. The dimensions of this bilayer are 45 × 45 nm, and the height of this system, including the two solvent layers at either side of the membrane, is 17 nm. In total, system A comprises approximately 300,000 CG atoms. The second simulation system, or system B, includes two ATP synthase dimers embedded in a much larger lipid bilayer, comprising 68,000 POPC molecules. The structural models for the ATP synthase dimers are based on cryo-EM data and X-ray structures, as described previously (Davies et al., 2012). The dimensions of the lipid bilayer are 85 × 254 nm, and the height of the system is 33 nm. In total, this second simulation system amounts to approximately 6 million CG atoms.

All molecular dynamics simulations we carried out with GROMACS 4.5.5 (Hess et al., 2008) and PLUMED (Bonomi et al., 2009). The integration time-step was 10 fs. The simulations were carried out at 300 K and 1 bar. A velocity-rescaling algorithm with time-constant 0.3 ps was used to preserve the system temperature. The pressure was sustained semi-isotropically using the Berendsen method, with a time-constant of 3.0 ps. Non-bonded interactions were calculated using a Coulomb potential, with relative dielectric constant ε = 15, and with a Lennard-Jones potential; a smooth shifting function was used so that both potentials are zero at 1.2 nm.

### Calculation of the bending modulus of the lipid bilayer

A molecular dynamics simulation of system A was used to estimate the characteristic bending modulus *k*_c_ of a symmetric POPC membrane, as represented by the MARTINI 2.1 force field. The bending modulus can be derived from the undulation spectrum of the membrane, which in turn can be calculated from a molecular dynamics trajectory. Specifically, in the low-frequency regime, the amplitudes of the different fluctuation modes of the membrane, *I* (*q*), fulfill the relationship (Helfrich, 1973; Marrink et al., 2004):

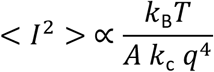

where < > denotes a time-average, *A* is the membrane surface area, *q* is the mode wavenumber, *T* the temperature and *k*_B_ the Boltzmann constant. A trajectory of 275 ns was calculated for this purpose. The undulation spectrum was then obtained from the last 150 ns of this trajectory. Specifically, a Fourier transform of the *z* positions of the phosphate groups in each leaflet was calculated for each trajectory snapshot, and the resulting amplitudes were averaged over the trajectory. The value of *k*_c_ was then estimated by fitting the function shown above to the calculated spectrum in the low-frequency regime.

### Potential-of-mean-force calculation

To determine whether the membrane alone can drive the association of two ATP synthase dimers, simulations of system B were carried out to calculate the change in free-energy as a function of the distance between the two dimers, *r*. This one-dimensional free-energy profile, or potential-of-mean-force, was calculated using the umbrella-sampling method. The total simulation time was 15 microseconds. Specifically, 200 independent simulations were carried out spanning a range of *r* values from 2.5 to 50 nm, in increments of 0.25 nm (additional simulations were introduced in smaller increments in some cases to guarantee overlap of the calculated probability distributions). In each simulation, the value of *r* was maintained in the vicinity of the target using a harmonic restraining potential of force constant 95.6 kcal/mol nm^−2^. To simplify the calculation, the two dimers were kept at all times approximately parallel to each other. To do so, we monitored the distances between the centers-of-mass of opposing c-rings, and applied an additional restraint that minimizes their difference, using a harmonic potential of force constant 28.7 kcal/mol nm^−2^. The starting configurations for this series of simulations were produced sequentially, i.e. for a given simulation, the initial snapshot was extracted from the preceding simulation along *d*. Each simulation comprised a relaxation stage of 37.5 ns, followed by a production phase of 70 ns. Snapshots were collected every 1 ps of simulation time, i.e. over 15 million snapshots were analyzed. The free-energy profile was calculated using the DHAM algorithm (Rosta and Hummer, 2015). To estimate the statistical error, the simulation data was split into two sets, and the mean difference of the corresponding PMF profiles was calculated.

### Membrane curvature analysis

The elastic deformation of the membrane induced by the two ATP synthase dimers was quantified as described previously (Davies et al., 2012). Briefly, for a given position in the plane of the membrane (*x*, *y*) and a given simulation snapshot, we evaluate the instantaneous membrane curvature in the *x* and *y* directions by calculating the radius of the circle that best fits the membrane surface at that position, in either direction. The reciprocal of this radius value is the instantaneous local curvature, *c*_x_ or *c*_y_. The mean curvature *c* is the sum of these two local curvatures (Marsh, 2006), time-averaged over the simulation length i.e., < *c* > = < (*c*_x_ + *c*_y_) >. To calculate the membrane curvature for each snapshot of the simulation, the phosphate groups of the lipid molecules were mapped on a two-dimensional grid in the (*x*, *y*) plane (0.5 nm grid-point spacing), based on their proximity to each grid point, using a cut-off distance of 9 Å. The circular fitting was then carried out at each grid point of the average surface following the method of Forbes (Forbes, 1989).

## Results

### ATP synthase dimers are driven together by a long-ranged membrane-induced force

To evaluate whether ATP synthases are organized by a membrane-mediated attractive force, we simulated the association of two ATP synthase dimers embedded in a phospholipid membrane of dimensions 250 nm by 85 nm, surrounded by solvent (**Fig. 2**). The number of particles in this simulation model was approximately 6 million. A series of 200 molecular-dynamics trajectories were then calculated to probe the propensity of the dimers to move closer together or further apart as the distance between them, *r*, was gradually varied (Methods). These simulations used the umbrella-sampling method, totaling 15 microseconds of sampling time. From a global analysis of these trajectories, we deduced the gain or loss in free-energy resulting from the change in the dimer-to-dimer distance, i.e. the potential-of-mean-force *W*(*r*).

**Figure 2.**
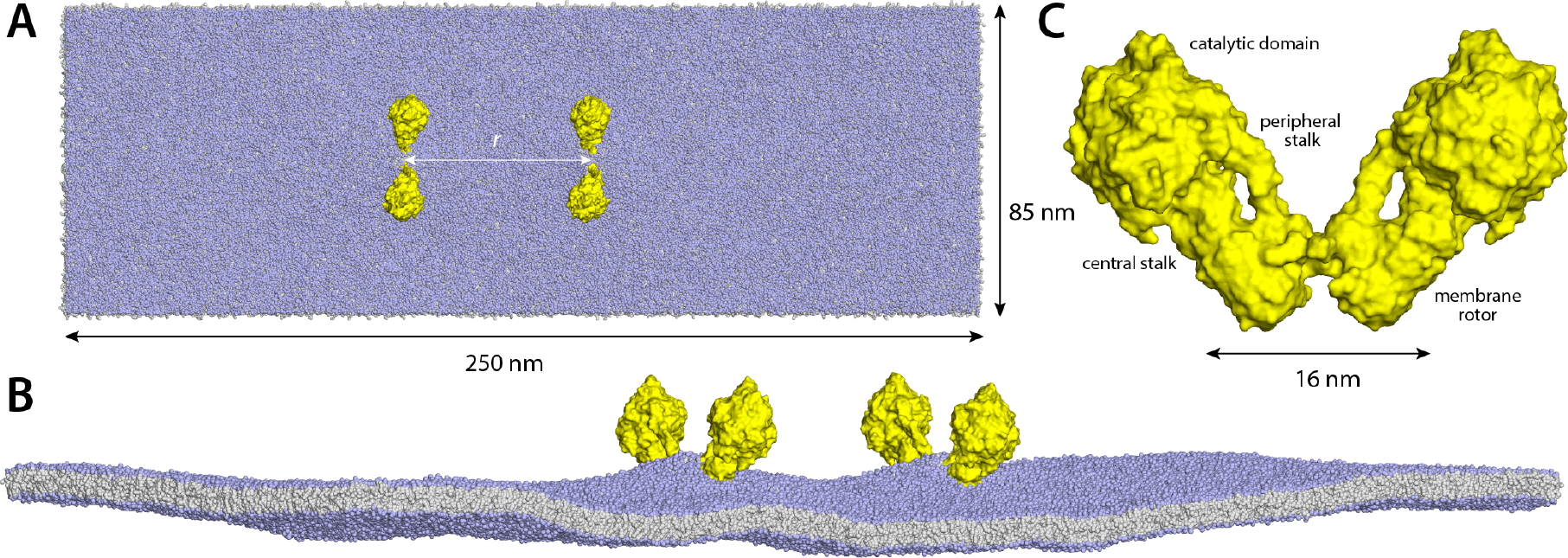
Molecular-simulation system comprising two ATP synthase dimers in a model lipid bilayer. (**A,B**) Views of the system along the perpendicular to the membrane and along the membrane plane, respectively. The calculation is based on a coarse-grained (CG) representation of all molecular components (Methods). The ATP synthase dimers are shown in yellow, in a surface representation. Lipid molecules are represented with spheres, with the head-groups (choline and phosphate) in purple and the acyl chains in grey. The solvent is omitted for clarity. The number of particles in the system is approximately 6 million. (**C**) Close-up view of an ATP synthase dimer, seen along the membrane plane. The structure is based on earlier cryo-EM studies. In the dimer, the monomers are approximately at a right angle.

The results obtained very clearly demonstrate that the two dimers are driven together by a strong, long-ranged force induced by the membrane (**Fig. 3**). Specifically, the calculated potential-of-mean-force shows that this effective interaction begins to act at distances smaller than 40 nm (center-to-center), when the membrane deformations induced by each dimer begin to coalesce, gradually forming a ridge in between the dimers (**Fig. 3A**). The free-energy of the molecular system decreases steadily as the dimers come closer together, reaching a minimum at 11-13 nm (**Fig. 3B**). Up to this point, the dimers are never in contact, and direct physical interactions between them are negligible. Beyond this point, however, the free energy rises quickly as the dimers begin to interact, and ultimately their steric repulsion becomes dominant. Thus, these large-scale free-energy calculations show the membrane alone brings the ATP synthase dimers together, up to an optimal non-contact distance where the two dimers form a membrane ridge, flat in cross section (**Fig. 1**).

**Figure 3.**
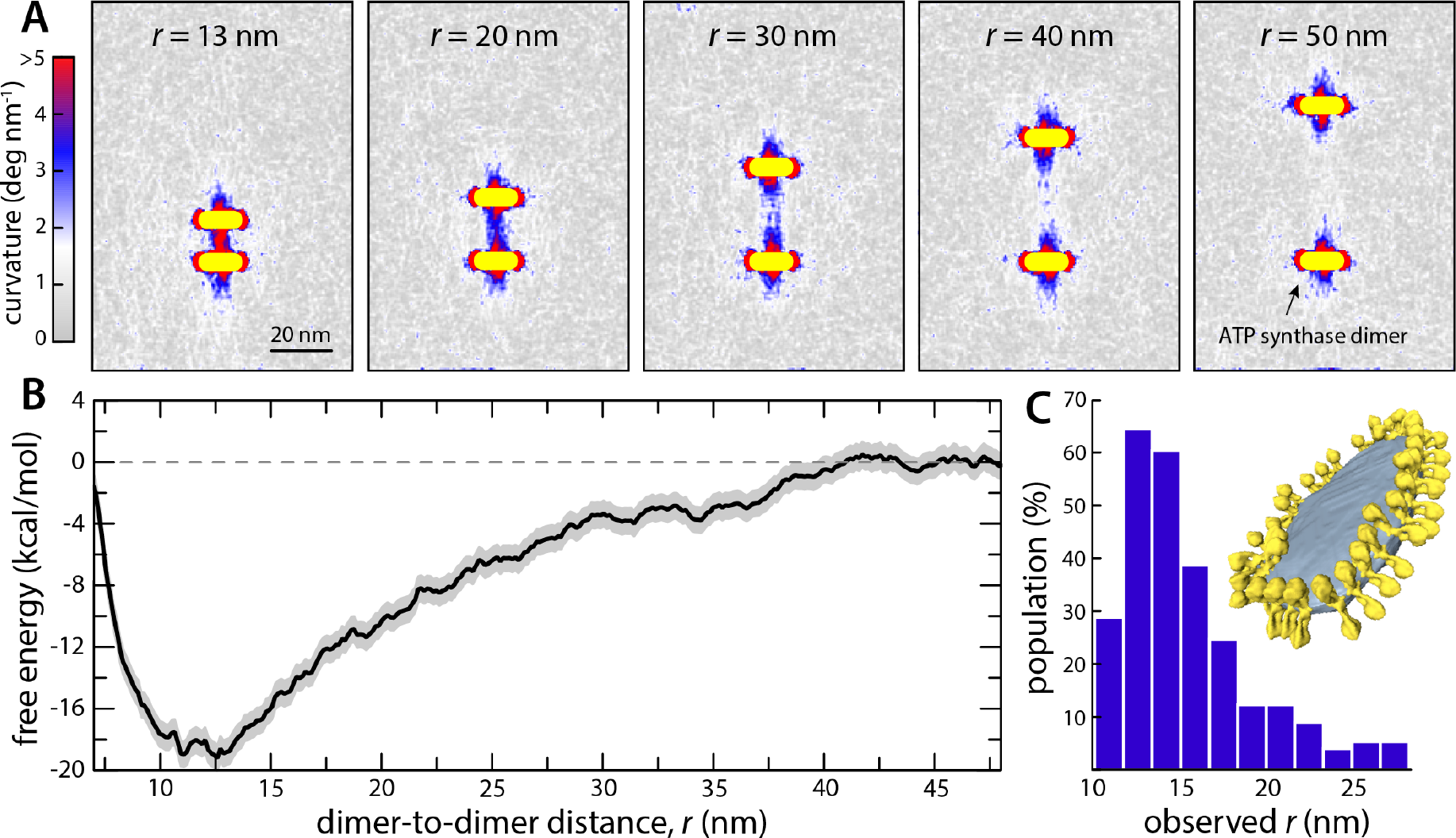
Energetics of association of two ATP synthase dimers, driven by the membrane. (**A**) Deformation of the membrane induced by two ATP synthase dimers, for different values of the dimer-to-dimer distance. The deformation is quantified by the position-dependent curvature of the membrane (Methods) along the two directions defining the membrane plane. (**B**) Potential-of-mean-force as a function of the distance between the two dimers, calculated using umbrella-sampling MD simulations, totalling 15 microseconds (Methods). The sampling error is indicated with a grey band along the free-energy profile. (**C**) Distribution of dimer-to-dimer distances deduced from analysis of published cryo-EM tomograms of fragmented mitochondrial cristae (inset figure adapted from (Davies et al., 2012)).

### The most favorable dimer-to-dimer distance matches experimental observations

Analysis of electron cryo-tomograms of mitochondrial membranes from *Saccharomyces cerevisiae* (Davies et al., 2012) show that the typical distances between ATP synthase dimers observed *in situ* are highly consistent with those predicted by the theoretical model. Specifically, the most frequently observed distance between adjacent dimers in a row is 12 nm (**Fig. 3C**). Thus, the membrane-mediated attractive force quantified in the computational model appears to optimally explain the organization observed physiologically.

The magnitude of the calculated free-energy minimum is also worth noting. To translate this value into a “bound-state” probability, one must integrate the one-dimensional profile *W*(*r*) to deduce a two-dimensional dissociation constant:

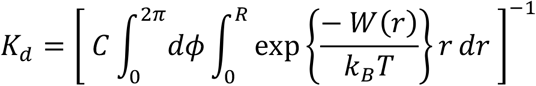

where *k*_*B*_ is the Boltzmann constant, *T* the temperature and *R* is a threshold in the dimer-to-dimer distance differentiating bound and unbound states. Note that the units of *K*_d_ in the above expression are molecule/nm^2^. While this expression is general, the quantity *C* is specific to our case, as it is a correction for the fact that throughout the calculation of *W*(*r*) the dimers are kept approximately parallel, for computational convenience (Methods). To a first approximation, *C* can be estimated as Δθ/2π, where Δθ is the rotational fluctuation of one dimer relative to the other, around the membrane perpendicular, in the associated state (note the rotational freedom of the unbound dimer is 2π, and not 8π^2^, as in solution, on account of the orientational restrictions imposed on the dimer by the membrane itself). The calculated values of *K*_*d*_ and *C* from the simulated data are ~3 × 10^−15^ nm^−2^ and ~0.03, respectively. Finally, the bound-state probability is:

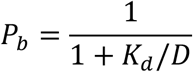

where *D* is the area-density of dimers in the membrane. For example, the system depicted in Fig. 2 approximately corresponds to a mole-fraction of 5 × 10^−5^ (one dimer per 20,000 lipids) which translates into *D* ~ 7 × 10^−5^ nm^−2^ (for an area-per-lipid of ~70 Å^2^). At this density, the population of the dissociated state would be virtually zero; specifically, 1 − P_b_ ~ 10^−10^. Although biological densities might be significantly lower, and the multi-dimer free-energy profile describing the assembly of a row would be shallower than that for two dimers only, this analysis clearly indicates that isolated dimers will be extremely rare in membranes.

### Model lipid bilayer reproduces experimental bending module

The question posed in this study cannot be quantitatively evaluated using a fully atomistic representation of the molecular system. Hence, we have resorted to a coarse-grained representation, specifically that implemented in MARTINI force field. Whether or not this widely-used approach is sufficiently accurate to study close-ranged interactions between membrane proteins has been recently questioned (Javanainen et al., 2017). The process we described does not entail such protein-protein interactions and therefore MARTINI appears a valid approximation in our view. Nonetheless, and given that the process we describe is driven by the membrane’s resistance to deformation, we asked whether MARTINI provides a reasonably accurate description of the elastic properties of the specific model lipid membrane considered in this study. We therefore carried out an MD simulation of a protein-free POPC bilayer (~5,800 lipid molecules), and calculated its bending modulus from analysis of its undulatory spectrum (Methods). As shown in **Fig. 4**, the intensities of the low-frequency modes < *I*^2^ > decrease linearly as a function of the forth power of the wavelength, *q*, as predicted from elastic theory (Helfrich, 1973; Marrink et al., 2004). The bending modulus *k*_*c*_ can be derived from the slope of the line fitting the calculated < *I*^2^ > as a function of *q* in a logarithmic scale. The calculated value of *k*_*c*_ from this analysis is 1.2 × 10^−19^ J, which is in good agreement with the value experimentally determined for POPC, namely ~1 × 10^−19^ J (Marsh, 2006). The coarse-grained MARTINI representation therefore appears to be a valid approach to evaluate the association process considered this study, notwithstanding the simplifications inherent to this force-field and its reported limitations for other types of processes controlled by direct protein-protein interactions (Javanainen et al., 2017).

**Figure 4.**
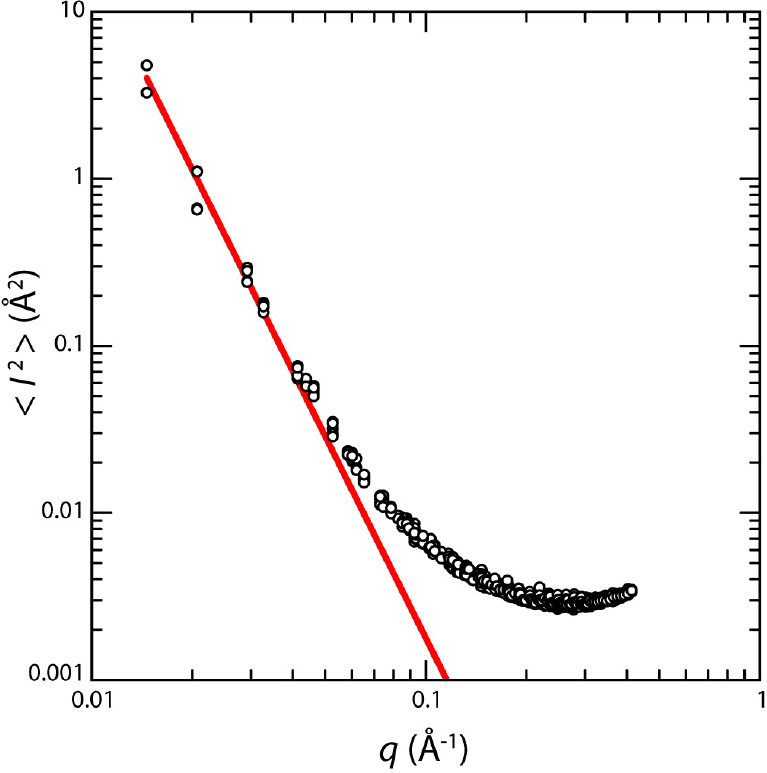
Bending modulus of the model phospholipid membrane used in the simulations. The calculation was carried out for a membrane patch without any protein (see Methods). The calculated value deduced from the slope of the linear fit (red line) is 1.2 × 10^−19^ J, in good agreement with the value experimentally determined, namely ~1 × 10^−19^ J (Marsh, 2006).

## Discussion

### Self-association requires dimerization but not direct dimer-dimer contacts

It has long been assumed that mitochondrial ATP synthase dimers assemble into rows as a result of direct protein-protein interactions between dimers (Minauro-Sanmiguel et al., 2005; Dudkina et al., 2006; Thomas et al., 2008; Couoh-Cardel et al., 2010). This notion stems in part from analysis of two-dimensional EM images of detergent-solubilized dimers, which appeared to reveal two populations differing in the relative angle between monomers (Dudkina et al., 2006). This result seemed to indicate the possibility of conformational adjustment upon direct interaction between dimers. However, subsequent tomographic analyses of mitochondrial cristae have shown that the conformation of the dimer *in situ* is invariant for a given species (Davies et al., 2011; Davies et al., 2012). Moreover, this tomographic data is in agreement with three-dimensional structures determined through single-particle cryo-EM (Hahn et al., 2016). A likely explanation for this discrepancy is that the reported structural heterogeneity of the dimer (Minauro-Sanmiguel et al., 2005; Dudkina et al., 2006; Thomas et al., 2008; Couoh-Cardel et al., 2010) results from different two-dimensional projections of the same conformation. Accordingly, in our simulations we study an invariant structure of the dimer throughout the range of distances considered.

The concept that dimer-to-dimer contacts occur also originates in biochemical and mutational studies indicating that subunit 4 (or *b* in bovine), in the peripheral stalk, and subunits *e* and *g*, in the transmembrane region, are involved in oligomerisation. These subunits were proposed to mediate two different dimerization interfaces, thereby generating the arrangement of the dimers observed in mitochondria (Everard-Gigot et al., 2005). Subunit 4 was specifically proposed to form dimer-to-dimer contacts through its soluble region (Spannagel et al., 1998). However, the more recently obtained high-resolution structures of the yeast ATP synthase dimer make very clear that subunits *e* and *g*, and the N-terminal transmembrane helix of subunit 4 are located at the interface between the two monomers of a dimer (Hahn et al., 2016; Guo et al., 2017). Indeed, the interaction between these subunits is the reason for the striking V-shape of the complex. The cryo-EM structures of the ATP synthase dimer also make clear that the peripheral stalks project away from each other, and thus the proposed interaction between the soluble regions of two subunits 4 cannot take place either within a dimer or between dimers. A probable explanation for this discrepancy is that the abovementioned cross-linking experiments were carried out with samples lacking subunit *e* and *g*, i.e. for monomeric ATP synthases. It is thus questionable whether these interactions are biologically relevant.

Subunit *e* has also been suggested to be involved in stabilising the ATP synthase dimer rows, through the formation of a coiled-coil by the C-terminal helix (Spannagel et al., 1998; Brunner et al., 2002). However, this putative interaction also appears to be incompatible with the cryo-EM structures of the yeast enzyme, which clearly show this C-terminal helix projecting straight down into the inter-cristae space (Hahn et al., 2016; Guo et al., 2017). The fact that this element could be discerned even at 7-Å resolution suggests that it is relatively rigid and thus unlikely to adopt an entirely different conformation. In summary, although the notion of direct dimer-to-dimer interactions is intuitive, the supporting biochemical evidence is, in our view, unconvincing, as it is incompatible with structural data more recently obtained through both high-resolution single-particle analyses and cryo-tomography of mitochondrial membranes. We therefore posit that subunits *e*, *g* and 4*(b)* are important for the formation of the ATP synthase rows not because they mediate interactions between adjacent dimers, but simply because they are required for the dimerization of the enzyme, and more specifically, its V-shaped configuration. Accordingly, the preferred distance between dimers observed in both experiment and computation (around 12 nm) is much too large for a direct contact, or even a direct physical interaction (e.g. electrostatics).

### Potential experimental validation and expected outcomes

The computational analysis presented here generates two clear-cut experimentally testable predictions. First, that ATP synthase dimers will self-associate into rows without requiring any other protein or a specific membrane component, ultimately creating a membrane ridge along the length of the row. Second, that the driving force for the formation of dimer rows originates in the membrane deformation imposed by the V-like shape of the dimers. To test the first hypothesis, purified ATP synthase dimers could be reconstituted into tethered lipid bilayers or synthetic liposomes; the distribution of these dimers in these membranes could be then analyzed using high-resolution atomic-force microscopy or electron cryo-tomography. To test the second prediction, the same experiment could be conducted for ATP synthase monomers, e.g. constructs lacking the dimer-specific subunits *e* and *g* (Davies et al., 2012). In the first experiment, we expect that the dimers will self-associate into long rows; indeed, the magnitude of the attractive force computed here suggests that isolated dimers will be seen rarely or not at all. The second experiment, by contrast, should show no evidence of association or membrane organization.

### The ATP synthase as a membrane-bending protein crucial for cristae morphology

All cellular membranes undergo constant shape changes. Gentle curvatures can occur through the asymmetric distribution of lipids, whereas acute curvatures require specialized proteins, such as clathrin or BAR, which coat the membrane surface and alter its morphology through the formation of a large oligomeric structures (McMahon and Boucrot, 2015). Mitochondrial cristae are formed from the repeated infolding of the inner membrane. This process is driven by lipid biosynthesis, and the resulting increase in the surface area of the membrane. As this growth occurs in a confined space i.e. within the outer membrane, the inner membrane buckles inwards. However, membrane growth alone does not explain the morphology of cristae. Mathematical modelling of this growth process shows that although multiple buckling events might occur initially, fusion events would eventually lead to a several, large vesicle-like compartments (Renken et al., 2002; Kahraman et al., 2012). Mitochondrial ATP synthases are unlike most other membrane-bending proteins in that they are integral transmembrane complexes whose primary biological function, *individually*, is not membrane remodelling. Nonetheless, we propose the biological role of the ATP synthase *dimers* is to prevent this kind of vesiculation, and to foster instead flattened lamellar-shaped cristae, or other types of geometry that provide order to the increasing surface area of the inner mitochondrial membrane. We envisage that early on in the development of the mitochondrion, rows of ATP synthase dimers form at multiple locations in the inner membrane, priming these sites for invagination. As the membrane grows, the inner membrane preferentially buckles at these primed locations, carrying the rows of dimers into the matrix. Due to the strong membrane curvature generated by the row, the invaginations would adopt a flattened-disk geometry, and the presence of ATP synthase dimer rows along the edges would prevent unregulated fusion or division of the inner mitochondrial membrane (**Fig. 5**).

**Figure 5.**
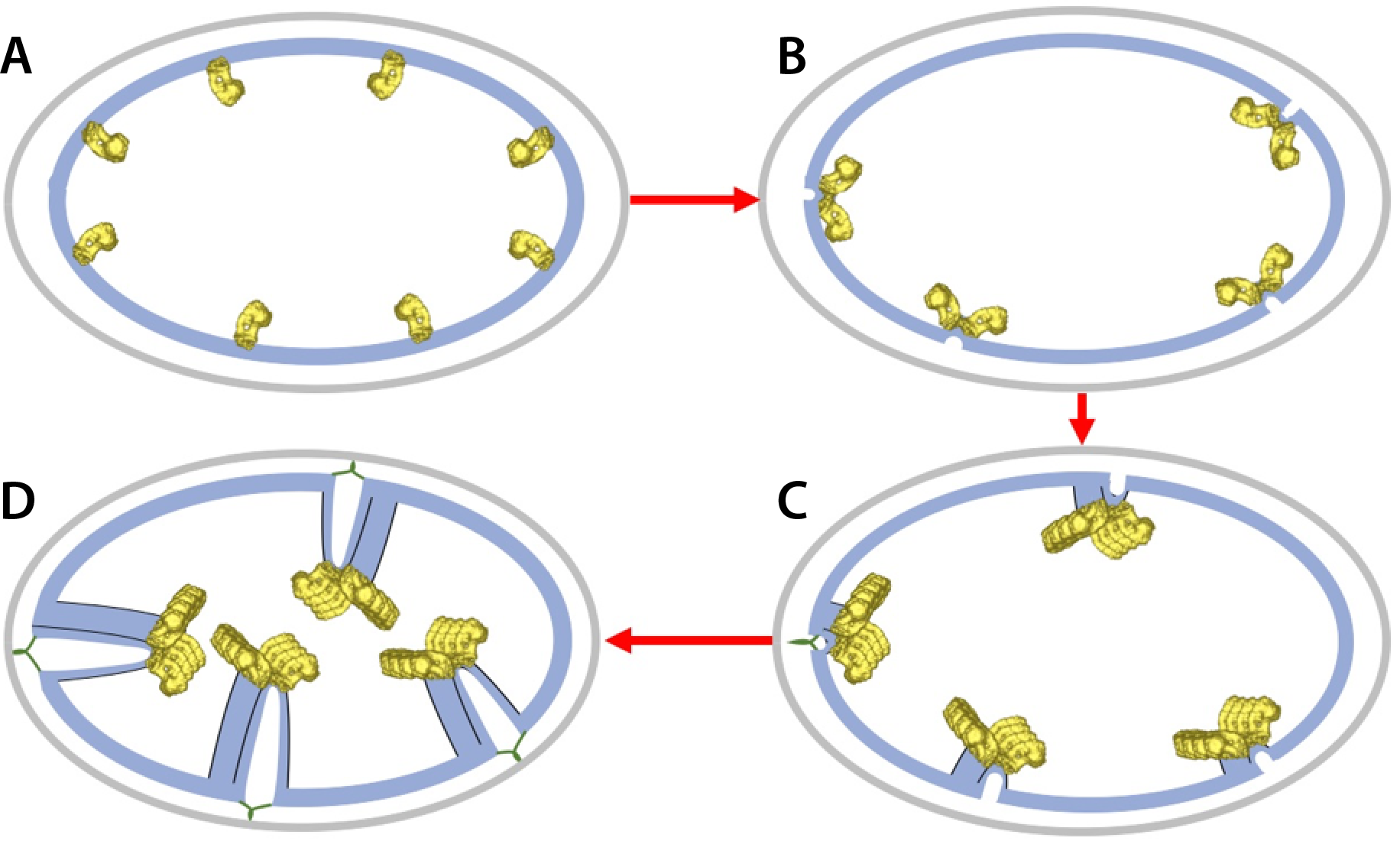
Proposed mechanism of cristae formation induced by the ATP synthase dimers. (**A**) ATP synthases (yellow) fold and assemble as monomers in the membrane. (**B**) As the ATP synthase monomers dimerise they cause a long-ranged deformation in the membrane which extends up to 40 nm away from the complex. (**C**) As the dimers encounter each other, the energy gained from reduced membrane curvature drives the dimers to self-assemble into rows. (**D**) The dimer rows form a macroscopic membrane ridge that primes the inner membrane to fold and as its surface area increases during mitochondrial development; the membrane invaginates exactly at the location of the dimer rows and cristae are generated. Inner membrane, light blue; outer membrane, grey.

## Conclusions

We have presented a molecular-simulation analysis designed to probe the factors that govern the membrane organization of mitochondrial ATP synthase dimers *in vivo*. The results presented explicitly demonstrate that there exists a force that drives the dimers together, side-by-side, to form or extend a row. This attractive force originates in the membrane deformation created by the structure of the isolated ATP synthase dimer. This deformation is energetically costly, but is gradually reduced as the dimers approach each other to form a ridge. The range of this effective attractive force is tens of nanometers, where no direct physical interactions between dimers occur. At the most favourable dimer-to-dimer distance, shown to be around 12 nm by both computations and experiments, protein-protein contacts cannot occur either. We therefore conclude that the formation of dimer rows is a process of spontaneous self-association, driven by the membrane itself. Importantly, this hypothesis is experimentally verifiable, for example through in vitro reconstitution assays of purified ATP synthase dimers and monomers. The mechanism we propose does not require a specific lipid composition, since all membranes resist deformation. It is essential, however, that the V-shape of the dimer is intact. We further posit that rows of ATP synthase dimers prime the inner mitochondrial membrane to develop cristae, and contribute to determining their shape and stability. Thus, these remarkable enzymes would not only produce most of the ATP consumed by the cell, but also help define the macroscopic morphology of the organelle in which they operate.

## Acknowledgements

This work was supported by the Max Planck Society (CA, KMD and JDFG); by the Cluster of Excellence “Macromolecular Complexes” of the Deutsche Forschungsgemeinschaft (KMD and JDFG); and by the Division of Intramural Research of the National Heart, Lung and Blood, National Institutes of Health (CA and JDFG). Computational resources were in part provided by the Biowulf HPC system at the National Institutes of Health, the Jülich Supercomputer Center, and PRACE. We thank L.R. Forrest, V. Leone and F. Marinelli for their comments on this manuscript.

